# *wMel* replacement of dengue-competent mosquitoes is robust to near-term climate change

**DOI:** 10.1101/2022.10.12.511962

**Authors:** Váleri N. Vásquez, Lara M. Kueppers, Gordana Rašić, John M. Marshall

## Abstract

Rising temperatures and increasing temperature variability are impacting the range and prevalence of mosquito-borne disease. A promising biocontrol technology replaces wild mosquitoes with those carrying the virus-blocking *Wolbachia* bacterium. Laboratory and field observations show that the most widely used strain, *wMel*, is adversely affected by heat stress. Here, we examine whether and how climate warming may impact *wMel*-based replacement. We integrate empirical data on the temperature sensitivity of *wMel* bacteria into a mechanistic model of population dynamics for the dengue vector *Aedes aegypti* and use CMIP5 climate projections and historical temperature records from Cairns, Australia to simulate vector control interventions. We show that higher mean temperatures are predicted to lower *wMel* infection frequency and that extended heatwaves have the potential to reverse the public health benefits of this intervention. Sensitivity analysis probing the thermal limits of *wMel* replacement reveal that, under existing projections, operational adaptations would be required for heatwaves lasting longer than two weeks. We conclude that this technology is expected to be robust to both the increased mean temperatures and heatwaves associated with near-term climate change in temperate regions. However, more rapid warming or tropical and inland regions that presently feature hotter baselines may challenge these tested limits, requiring further research.

## Introduction

Climate change is rapidly altering the global landscape of vector-borne disease.^1–3^ Numerous empirical studies have demonstrated that temperature influences both mosquito and pathogen traits, affecting the geographic range and prevalence of illnesses such as malaria, dengue, and Zika virus.^4–6^ Anthropogenic warming is expected to cause heterogenous impacts, with some regions seeing increases in seasonal suitability for the malarial *Anopheles* mosquito and others undergoing extensions of the environmental conditions favorable to the *Aedes* mosquito that transmits dengue and a range of other arboviruses. Temperate areas are projected to experience augmented periods of annual exposure to both vector groups.^7^

The emerging scientific understanding of disease risk under future climates underscores a need for improved vector control tools, particularly as insecticide efficacy wanes due to evolved resistance^8–10^. There has been a surge of innovation in control technologies over the past decade, including several successful field trials of a self-sustaining biocontrol approach called *Wolbachia*-based replacement.^11–13^ This technique, which transfects wild mosquitoes with a maternally-inherited endosymbiotic bacterium that is naturally-occurring in many arthropods, blocks carriers’ ability to transmit human pathogens.^14^ Multiple strains of *Wolbachia* bacteria, each bearing unique biological properties, have been transferred into various *Aedes* species (*Ae. albopictus, Ae. polynesiensis*, and *Ae. aegypti*) in public health intervention trials spanning Latin America, Asia, and Oceania.^15^ Most *Wolbachia* replacement efforts trialed to date have used the *wMel* strain.^16^

*wMel* is a naturally occurring symbiont of *Drosophila melanogaster*. In transfected mosquitoes it reliably causes stable maternal transmission and blocks replication of pathogenic viruses while incurring a relatively low fitness cost compared to alternative *Wolbachia* strains.^17^ Under baseline laboratory conditions *wMel* demonstrates complete cytoplasmic incompatibility (CI), which renders crosses between infected males and uninfected females unviable, favoring the reproduction of *wMel* carriers and driving its own spread through a vector population. However, recent studies have shown that CI – a property vital to *wMel* utility in public health applications – is weakened by cyclical heat stress, including the daily and seasonal thermal fluctuations that can characterize field conditions.^18^ High temperatures have also been shown to decrease the hatch rate of *wMel*-infected eggs and reduce *wMel* density in adults, impairing maternal transmission and causing the infection to fall out of a laboratory population once a threshold of 35.0°C is reached.^19^

Using projections of future increases in the frequency, severity, and duration of heatwaves^20^ together with laboratory evidence of the negative impact that oscillating temperature regimes and thermal shocks have on *wMel* infection,^18,19^ we test our hypothesis that the hotter, more prevalent heat extremes and higher average temperatures brought by climate change could impede *wMel*-based public health interventions. We assess thermal effects on this technology in Cairns, Queensland, Australia – the location of the first successful field trials for *wMel* population replacement – using entomological endpoints, including the frequency of *wMel* infection and the magnitude of wildtype suppression in a simulated *Ae. aegypti* population. The computational experiments developed to probe our hypothesis explore the compounding interaction of warming trends, *wMel’s* intrinsic biological thresholds and extrinsic fitness costs, and uncertainties about the functional form of temperature impacts on that symbiont.

## Results

### Increased Average Temperature

Results demonstrate that *wMel*-based population replacement is generally robust to projected regional warming trends out to mid-century. Outcomes were quantified using Suppression Efficacy Score (SES)^21^ – an index gauging the proportion between ideal efficacy (complete eradication) and observed result – as well as a new metric, Replacement Efficacy Score (RES), that was formulated for this work and likewise evaluated on a 0 to 100 scale relative to ideal efficacy (fixation). The RES score summarizes the success with which the standing population is substituted by *wMel* infected individuals (see online Methods). Over all scenarios, SES and RES scores remained above 76 and 81, respectively (Fig.1).

**Figure 1:**
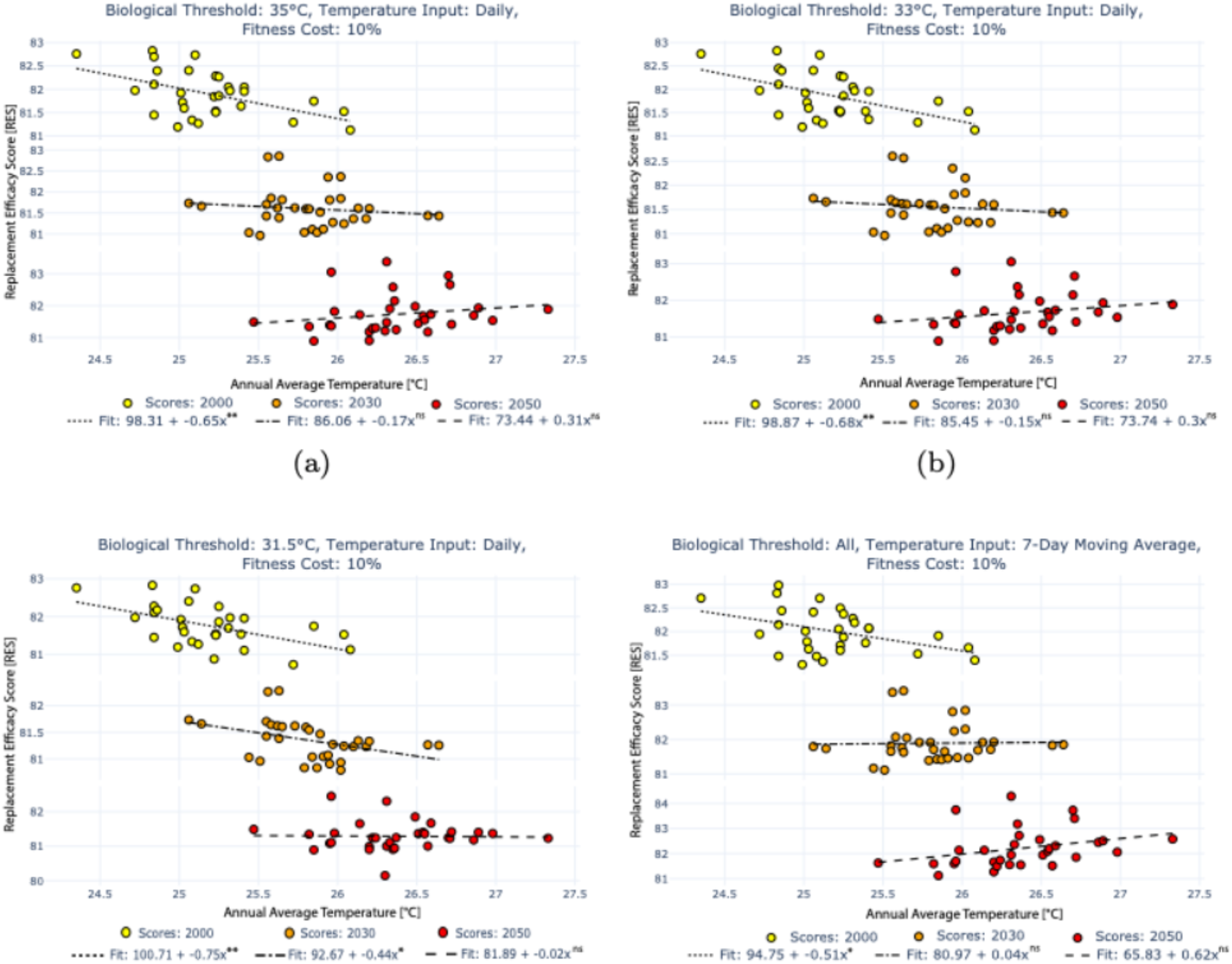
Increases in average temperature impact the replacement efficacy of Wolbachia-based interventions. Each circle marker corresponds to a Replacement Efficacy Score for the intervention conducted in an individual year, with colors corresponding to the temperature regime (historical, 2030 projected, 2050 projected). Trend line fits are noted in each panel legend. Panels A - C reflect results from the sensitivity analysis for unique thermal thresholds and daily temperature inputs to the wMel dynamic model. Panel D displays results when seven-day moving averages are used as temperature inputs to the wMel model; all three thermal thresholds returned equivalent output under this assumption.

We examined the consequences of two thermally sensitive biological mechanisms – CI and maternal inheritance – failing completely when daily average temperature reached the 35.0°C threshold which has been empirically shown to drop *wMel* titer to zero in exposed *Ae. aegypti* eggs. We also conducted a sensitivity analysis to evaluate the possibility that diminished *wMel* presence at lower temperatures, namely 33.0°C and 31.5°C, may have the same effect. For interventions conducted under average daily temperatures across both historical (1990-2019) and projected future (2024-2039, 2044-2059) years, RES scores and their correlation with temperature were nearly identical between 35.0°C and 33.0°C threshold scenarios (Fig.1a,b). The lowest threshold, 31.5°C, produced outcomes that were significantly more sensitive to temperature than those generated by the 35.0°C and 33.0°C alternatives (Fig.1c). The same held true for SES scores (Section 3 in Supplementary Information (SI)).

Because the data used to develop *wMel* temperature-responsive dynamics were from laboratory studies conducted at a temporal resolution of one week, we performed a sensitivity analysis to investigate the effect of altering the functional form of temperature inputs. In one set of simulations temperature inputs to the mechanistic model of *wMel* temperature sensitivity realized a daily effect, reflecting the assumption that the same impact recorded after one week of exposure would also occur after one day. In a second set of simulations, a seven-day moving average constituted the temperature inputs to the *wMel* model, emulating the empirical data generating process. The simulations driven by moving averages resulted in equivalent output under all three threshold values (35.0°C, 33.0°C, 31.5°C) for thermally responsive CI and maternal inheritance (Fig.1c,d). These runs also generated a larger year-on-year difference between 2030 and 2050 RES scores and trends, as compared to results driven by daily temperature inputs. A third sensitivity analysis examined how fitness cost on *wMel* carriers affected outcomes. Because results were minimally influenced by fitness cost, all results featured in the main text assume a 10% cost; results for 0% and 20% cost are reported in Section 3 of the SI.

The sensitivity of *wMel* replacement to increasing temperature diminishes in future, warmer decades. **Figure 1** shows the RES scores for interventions conducted under average daily temperatures across both historical (1990-2019) and future (2024-2039, 2044-2059) years, with the latter encompassing both RCP 4.5 and 8.5 scenarios. Under the historical regime, assuming daily inputs to the *wMel* model and using the thermal threshold of 35°C, temperature as a linear covariate accounts for approximately 25% (R^2^ = 0.2540, p < 0.0045) of the variation in the model fit (y = 73.44 - 0.31x), while the slope of that fit indicates that each degree of temperature increase lowers the RES score by 0.31. While this demonstrates a strong negative correlation (*ρ* = −0.5041), it leaves 75% of variation due to non-temperature related factors in this biological system. However, any relationship with temperature disappears in the 2030 and 2050 regimes. Holding the thermal threshold and daily input assumptions constant during model runs representative of 2024-2039, the effect of annual average temperature on RES score – assuming the highest threshold of 35°C – is no longer significant (R^2^ = 0.0157, p < 0.4948). This is also the case in the years 2044-2059 (R^2^ = 0.0389, p < 0.2788).

Changing the functional form of temperature inputs such that the *wMel* model is driven by a seven-day moving average yields the same reduction in future temperature sensitivity. As before, there is a negative relationship between temperature and RES score during the historical period (y = 94.75 – 0.51x), however, annual temperature only accounts for 17.2% (R^2^ = 0.17219, p < 0.02259) of the variation in that fit. In the linear model fit for the 2030s and 2050s, the correlation is absent (2030s: R^2^ = 0.00048, p < 0.90467, 2050s: R^2^ = 0.3110, p < 0.08316). The chief distinction from the model with daily temperature inputs to *wMel* dynamics, aside from having identical outcomes across all three thermal thresholds, is that the absolute magnitude of the slope in the linear model fit is invariably larger. This is sensible, given that the moving average approach extends the effect of a single hot day for an additional six days.

Because the RES score is a single value that summarizes success over the entire period of interest – in this study, one year – understanding the reason for this metric’s shifting year-on-year relationship with temperature requires a closer look at the population dynamics (Fig.2). Under the 2050 scenarios depicted in red, the number of *wMel-*infected females drops towards the end of the calendar year (approaching summer in the Southern Hemisphere). During this period, thresholds for temperature-sensitive *wMel* mechanisms (CI and maternal inheritance) are exceeded, leading to the reduction of the infected population. However, we also see that rising average temperatures during the coolest part of the year improve *wMel* replacement during that period. These midyear dynamics are strong enough to mask the decrease in infected females that occurs in later months when using a summary metric.

### Longer, More Severe, and More Frequent Heatwaves

While *wMel*-based population replacement is negatively affected by the regional temperature variability projected to occur under climate change, it remains robust to current predictions of future heatwaves in temperate Cairns. For a full description of the methodology used to construct the heatwave timeseries, see Methods and Section 1 of the SI. As with the average temperature change experiments, we conducted sensitivity analyses over the range of thermal thresholds, functional form of temperature inputs, and fitness costs.

Simulations employing daily temperature inputs see a drop in infection frequency across all thermal thresholds (35.0°C, 33.0°C, 31.5°C) for select years (Fig.3). This holds true for all assumptions of fitness cost. In the most severe cases, the proportion of the population carrying *wMel* falls to 62% when assuming a 10% fitness cost. Notably, this remains well above what is considered a critical threshold (20-30%) for achieving *wMel* fixation (100% infection frequency).^22,23^ When simulation dynamics were driven by seven-day moving average temperature inputs, *wMel* infection frequency rose faster than in the results obtained using daily inputs, and in no instance did frequency fall after reaching fixation.

These results are put into population-level perspective in Fig. 4, which shows the dynamics of *wMel*-infected and wildtype female mosquitoes for each biological threshold and temperature input assumption under future regimes. Dynamics occurring under heatwave years are overlaid on dynamics driven by the baseline temperatures used to construct those heatwave inputs, highlighting the delta generated by thermal shocks. The decline in *wMel* infection frequency and corresponding increase in vector-competent female mosquitoes indicates the potential public health impact of future temperature variability. Given the model assumption of daily temperature inputs, the population of *wMel* carriers is more than halved by December for select years in both the 33°C and 31.5°C scenarios. A decline is observable for those same years in the 35°C scenario. In all cases, this decrease corresponds to a rising proportion of vector-competent females.

Current empirical data suggests that *wMel* falls out of host organisms at temperatures equal to or exceeding a 35°C threshold.^19^ In the climate projections used as the basis of simulation, this threshold is reached or surpassed as a daily average in 2029^1^ and again in 2049^2^. While these examples of severe heat have visibly negative effects on simulated *wMel* interventions as shown in Figs.3 and 4, they were not sufficient to cause a complete removal of the pathogen-blocking bacterium from the mosquito population. However, recent years of observed temperature in Cairns have featured greater variability and hotter extremes than those currently used as baselines for climate projections for this study (Fig. 5).

Given the robustness of *wMel* to the climate scenarios used in this work and the potential for warmer or more variable potential futures in Cairns and elsewhere, we conducted a theoretical exploration of what might be required to remove the infection from a population by incorporating progressively more hot days in existing CMIP5 heatwave years. **Figure 6** features the impact of this thermal limit experiment on wildtype females and their *wMel*-carrying counterparts. See Methods for the approach used to construct these augmented temperature inputs.

Panels (a) and (b) of **Figure 6** illustrate the effect of including additional hot days in the existing timeseries of projected future temperatures. The surge in vector-competent mosquitoes is larger in the 2050s scenario than that of the 2030s due to the high thermal optima of the *Ae. aegypti* species. Further, three fewer “extra” days of heat are required in 2050 for the wildtype population to bounce back. However, even under the most extreme assumption of consecutive hot days (19 and 16 in the 2030s and 2050s scenarios, respectively), the population of *wMel*-infected mosquitoes persists at levels that while low are sufficient to permit a potential full rebound (Fig.6c,d).

## Discussion

There is a growing literature on the thermal biology of mosquitoes, with several studies illustrating its role in the impact of climate change on vector-borne disease. However, new public health intervention technologies may also be subject to the complexities of global warming. To date, no research has examined the potential effect of climate change on *Wolbachia*-basedcontrol methods, nor employed empirical data on the thermal sensitivity of *Wolbachia* to model the dynamics of this biocontrol tool. Further, published modeling work that explores the influence of temperature on mosquito-borne disease has focused on trends (increasing average temperatures) but not variability (the magnitude, duration, and frequency of heat waves).

Here, we demonstrate that rising average temperatures in Cairns, Australia under CMIP5-generated predictions may increase *wMel* replacement during winter months. However, we also show that future average temperatures in the summer may surpass the symbiont’s biological thresholds and cause the decline of the infected population. Recent field work in Nha Trang City, Vietnam supports our finding that such elevated temperature conditions can lead to a decrease in *wMel* infection frequency, with researchers noting that a heterogeneity of factors additional to temperature may influence *Wolbachia* establishment.^24^

Our results also exhibit the potential vulnerability of wMel-based population replacement to temperature variability under climate change, with simulations that account for hotter and more frequent thermal extremes showing diminished efficacy relative to results produced by simulations that strictly consider mean temperature. Heatwaves projected by CMIP5 scenarios lower *wMel*-infection frequencies; however, infection frequencies appear to recover under even the most severe existing projections of future heat shocks. In 2018 (11/26-11/28), a heatwave in Cairns, Australia where *wMel* infection had been established in the *Ae. aegypti* population since 2011 afforded scientists the opportunity to observe a natural experiment whose conclusions are consistent with our findings.^25^ At the end of November that year, recorded maximum temperatures reached 43.6°C. While *wMel* frequency did fall, it never dropped below 83% in juvenile mosquito stages, and recovered to nearly 100% by April 2019.

But during the period of this natural experiment in Cairns, the highest average daily temperature was only 33.75°C. To fully explore the potential edge case in which *wMel* is severely reduced or fully eliminated from a population following heat shocks, experimentation is needed to assess the effect of a daily average greater than or equal to 35°C, and to understand the interaction of temperature spikes with mosquito developmental stages on pathogen-blocking efficacy.^26^ Increasingly acute and prolonged heatwaves around the world highlight the need to test the thermal limits of *wMel* technology beyond the theoretical examples presented here, where approximately two to three weeks at or above 35°C caused an extreme reduction – but not a complete eradication – of the infected population. Such dynamics may be relevant to a proposed toxin-antidote gene drive based on the *Wolbachia* alleles responsible for CI.^27^

Further investigation is also required to discern the minimal exposure time at or above said threshold before critical mechanisms such as CI and maternal inheritance are impacted. As demonstrated by our computational experiments, finer time resolution concerning the effect of temperature on the hatch rate of *Wolbachia*-infected eggs will further improve scientific insight. Additional points of necessary research include the generational duration of deleterious effects following heat spikes. Finally, our results are necessarily a function of structural decisions made when developing the ODE model, including the choice of functions to define temperature-sensitive vital rate parameterizations.^28^ Therefore, analyses using alternative formulations may be merited as well.

Our modeling does not account for potential heat avoidance (heat-induced migration) of mosquitoes, ^29,30^ and thus does not consider the possibility that warming trends may drive *Wolbachia*-infected *Ae. aegypti* to seek cooler temperatures, maintaining their bacterial titer.^29^ Together with additional research into the connection between *Wolbachia* density and the mechanisms of CI,^31^ further scientific investigation of the temperature seeking behavior caused by this symbiont is needed.^15^ Importantly, dengue virus has also been shown to increase the thermal sensitivity of *Ae. aegypti* and coinfection with *Wolbachia* does not appear to furnish protection from dengue-induced thermotolerance effects.^32^

*Wolbachia* experts recognize that strain selection for a given intervention must account for local environmental realities.^15^ Recent studies have focused on deployment in extreme environments^33^ and the development of strains with increased phenotypic stability under heat stress.^34^ Even so, it is understood that *“*ideal Wolbachia strains for population replacement do not exist, requiring a trade-off between Wolbachia infection stability, host fitness costs, and pathogen blocking.”^15^ For decisionmakers to appropriately weigh the odds between *wMel* and its alternatives as we move into an uncertain and more variable climate future, computational and bench scientists should explicitly account for predictions of regional warming – including heat extremes – in their experimental designs. In the short term, temperature-conscious operational planning such as augmented deployment schedules may be able to compensate for the already-observed risk that infection frequencies drop or remain at low levels due to heat stress.

While the results presented here furnish reassurance that *wMel* replacement is a resilient technology in the face of climate change, they are predicated on temperature profiles constructed from pre-2006 baselines in a temperate area, and thus remain conceivably best-case scenarios. IPCC AR6 WG1 noted with high confidence that Australian land areas have warmed by 1.4°C in just over 100 years, and that annual temperature changes “have emerged above natural variability in all land regions [of Australasia].”^20^ Climate scientists also warn of a consistent trajectory of heatwave intensification in Australia, predicting a potential 85% increase in prevalence should warming reach 2.0°C.^35^ This highlights the inextricable link between climate mitigation policy and infectious disease management.^1^ Failure to curb carbon dioxide emissions will not only increase the proportion of the world exposed to deadly mosquito-borne illnesses due to rising temperatures, it may also undercut the utility of an otherwise demonstrably effective tool for vector control.

## Methods

### Climate Data

We obtained historical daily temperature data for Cairns, Queensland, Australia (latitude - 16.8736, longitude 145.7458) from the Global Historical Climatology Network (GHCN) database maintained by the National Centers for Environmental Information of the United States National Oceanic and Atmospheric Administration (NCEI-NOAA). We selected Cairns as the geographic location of interest because two towns in that region, Yorkeys Knob and Gordonvale, were the sites of the first successful field trial for population replacement using *wMel*; the reported trial outcomes therefore serve as experimental validation for simulations using local environmental variables.^11^ Additional *Wolbachia* releases have since taken place in the area, including in central Cairns.^36,37^

Projected future data for Cairns including both average temperatures and heatwaves under RCP scenarios 4.5 and 8.5 were developed from dynamically downscaled Coupled Model Intercomparison Project Phase 5 (CMIP5) global climate projections obtained from the Queensland Future Climate Dashboard (hereafter QFCD), which is maintained by the Science Division of the Queensland Department of Environment and Sciences (DES).^38,39^ These data are recorded in fractions of degrees of long-term change (°C) relative to the reference period (1986-2005). Because the available daily historical temperature data begin in 1990, we used the period 1990-2005 as the historical baseline and developed daily future climate scenarios using the anomaly method for the years 2024-2059. While years from 2060 onward exhibit more drastic temperature rise, we excluded these from the analysis to focus on outcomes most relevant to near-term technological decisions concerning *Wolbachia*-based public health interventions. Full details on the methodology used to construct future daily average temperatures and heatwaves are in Section 1 of the Supplementary Information (SI).

### Simulation Model of Population Dynamics

We developed a system of ordinary differential equations (ODE) to simulate mosquito population dynamics. The model, described in detail in Section 2 of the SI, portrays four life stages of the mosquito lifecycle: egg, larva, pupa, and adult.^40,41^ Adults emerging from the pupal stage are evenly divided between males and females; the population is assumed to be randomly mixing and all organisms within a particular life stage are equal with respect to their birth, death, and maturation rates. Larval stage mortality is modulated by logistic density dependence.^42^ Parameterization reflects the species-specific thermal biology of *Ae. aegypti*, which was selected for this study because it has been a primary host for *wMel* in the context of population replacement experiments including in the Cairns region. The model was initialized at equilibrium.

*Ae. aegypti* vital rates were calculated using temperature responsive functional forms unique to each life stage. These formulations and their parameterization were developed by Rossi, Ólivêr, and Massad to probe the effect of climate change on the *Ae. aegypti* lifecycle as well as its relevance to disease incidence.^43^ Assumptions used to adapt these equations to the underlying population model included employing literature-derived values for oviposition (63 eggs daily per female) and excluding the consideration of carrying capacity for egg laying.^44^ Field studies of *wMel* replacement interventions in the Cairns region informed the deployment schedule used for the simulations, where model releases paralleled reality to occur every seven timesteps from day 4 through day 63.^11^

### Biological Parameterization of wMel Temperature Sensitivity

*Wolbachia* carriers in this work are assumed to have temperature-responsive hatch rates, maternal inheritance rates, and cytoplasmic incompatibility levels. Empirical data generated from the cyclical heat exposure of *wMel*-infected *Ae. aegypti* mosquitoes was used to develop the equations that reflect thermal sensitivity.^19^ In these laboratory experiments, eggs were subjected to weekly temperature regimes that cycled daily between the maximum and minimum of a given 10°C range, with the coolest regime averaging 29°C and the warmest averaging 37°C. Data was also collected for eggs held at a consistent 26°C for the seven-day study period.

Extrapolating the results of week-long experiments to a dynamic model with daily timesteps requires assumptions about the unknown functional form of heat stress accumulation in wMel. To address this uncertainty and account for the lack of required temporal resolution in the empirical data generating process, we included a sensitivity analysis that assesses two theoretical alternatives: first, all temperature inputs to the thermally sensitive wMel equations were calibrated to consist of a seven-day moving average. This reflects an assumption that the physiological impact of heat stress is evenly distributed, with the effect of one hot day prolonged for a subsequent six days. In the second sensitivity analysis, all temperature inputs were permitted to enter as daily values. This reflects an assumption that the impact of heat stress is primarily immediate and does not necessarily result in a cumulative effect within the organism.

To formulate the biological model of wMel, we fit a function to the mean value of experimental replicates for each temperature regime. Equation 1 reflects the statistical best fit to egg hatch data. The hatch rate *T(C)* for infected eggs below 26°C was fixed at 0.91833 and above 36°C it was fixed at 0.0 in accordance with Ross et al (2019). Methods **Figure 1** compares Equation 1 dynamics with observed hatch rates.

**Figure 1:**
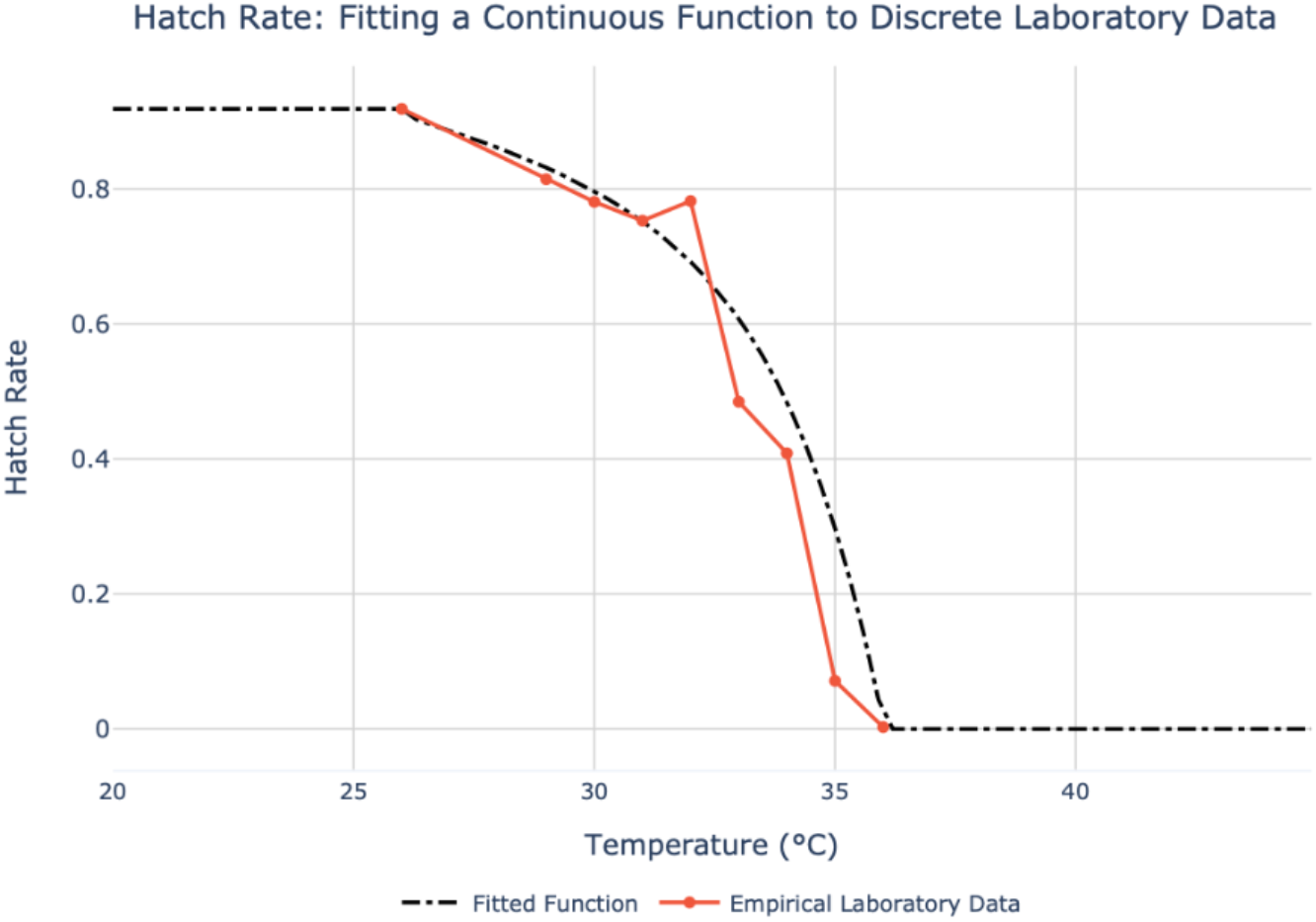
Fitted hatch rate function (Equation 1) plotted against empirical data underlying Figure 7a of Ross et al (2019), which recorded hatch rates for wMel-infected eggs under cyclical temperatures.

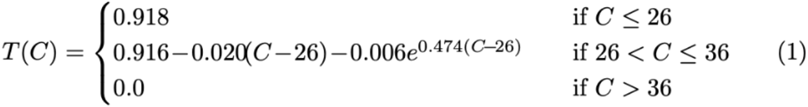

The temperature-sensitive status of *wMel* infection and consequent vertical transmission from mother to offspring was modelled using matrix operations informed by the data underpinning Figures 7B and 7C of Ross et al (2019), where above 35.0°C *wMel* inheritance, *I*, falls to zero and previously complete cytoplasmic incompatibility, *Y*, stops functioning entirely. These conditional relationships are formalized in Equations 2 and 3.

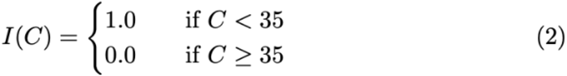

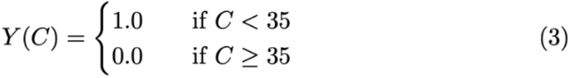

The conservative assumptions embodied by this approach, which rely on the simple presence or absence of *wMel* infection, are made in acknowledgment of the fact that the mappings between infection titer and the biological properties it confers are not yet well defined. The chosen method results in a likely overestimation of both *wMel* maternal transmission rates and cytoplasmic incompatibility at temperatures lower than the 35.0°C threshold. We therefore include additional sensitivity analyses varying this biological threshold, imposing it instead at 31.5°C and 33.0°C. Future work, including empirical experimentation to improve scientific understanding of the temperature sensitivities of *wMel* infection and mechanisms of cytoplasmic incompatibility, will enable refined mathematical estimations of these processes.

A range of fixed fitness costs was modelled to reflect heightened mortality in *wMel* carriers, drawing on empirically derived values in the literature. This sensitivity analysis tested costs that augmented adult mortality rates in *wMel*-infected mosquitoes by 0%, 10%, and 20%, with reported results reflecting use of the middle value (10%). Section 3 of the SI contains outcomes employing the upper and lower bounds of the fitness range. Field studies of *wMel* replacement interventions in the Cairns region are consistent with model dynamics, as illustrated in Methods **Figure 2** below comparing observed data with simulated results.

**Figure 2:**
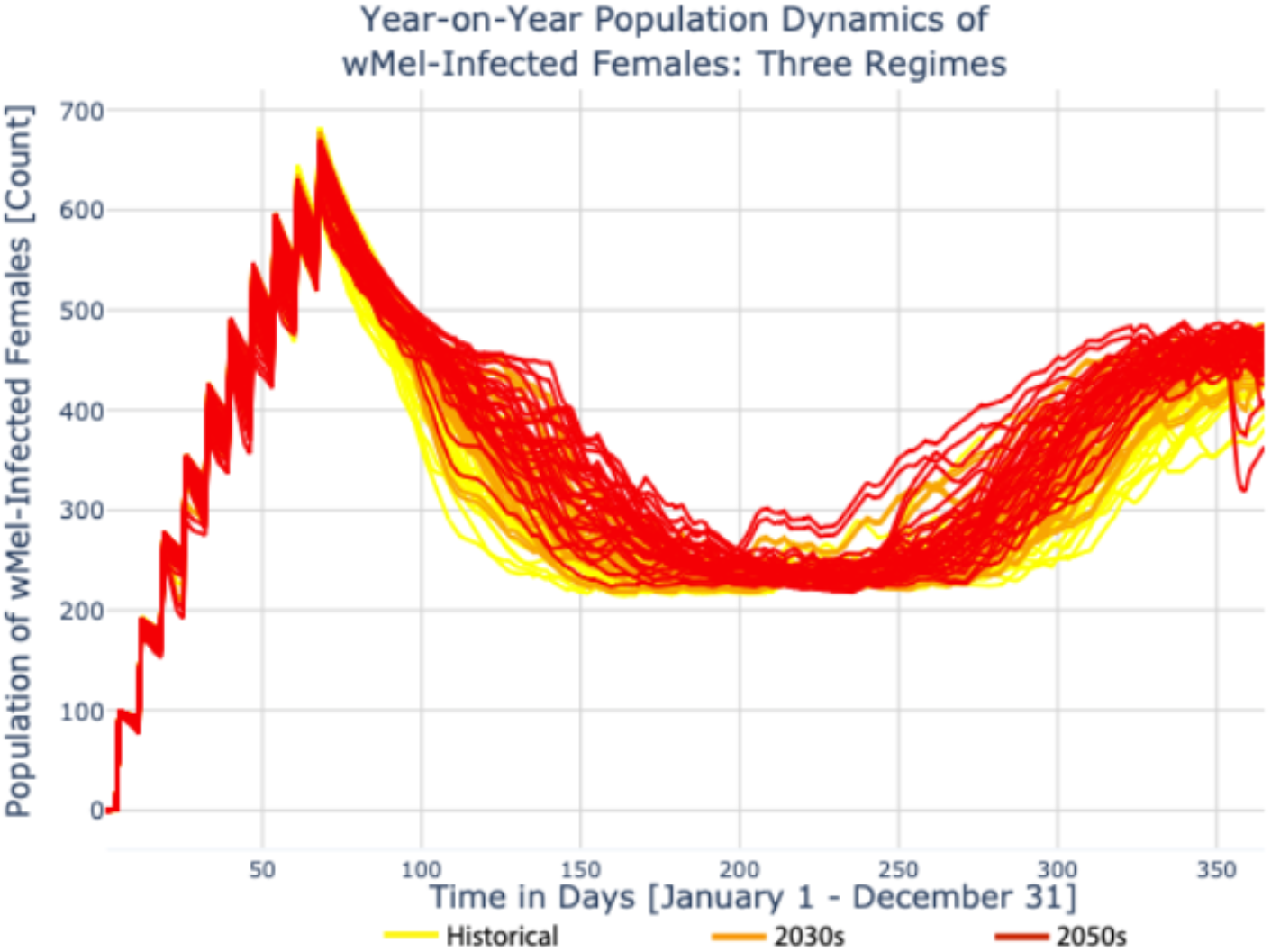
Effect of average temperature on the population dynamics of wMel-infected females lend insight to Replacement Efficacy Score results. Each Une corresponds to one simulated intervention year; each color corresponds to a different temperature regime (historical in yellow, 2030 in orange, 2050 in red).

**Figure 2:**
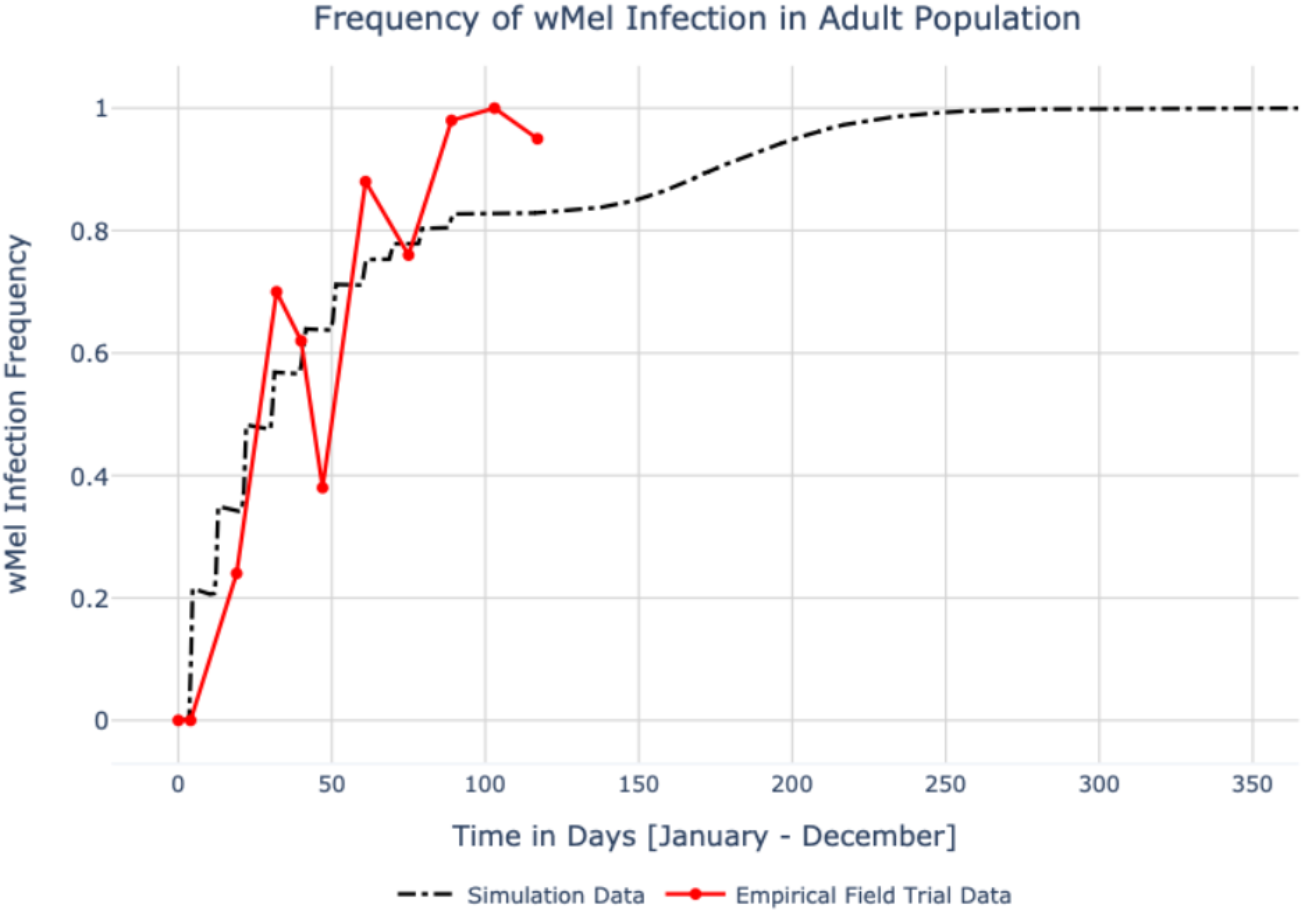
Empirical results from a 2011 field trial in Yorkeys Knob, Queensland, Australia support model dynamics. Here, a wMel-based replacement intervention with a release schedule based on that of the field trial is simulated using 2011 recorded temperatures, and results are compared with observed data. Figure 1A of Hoffman et al (2011) furnished the field observations used here. The final day of data collection in the field trial was April 27 (day 117); modelled results are shown through December 31 (day 365).

### Metrics Used to Assess and Compare Simulation Results

In addition to characterizing results using qualitative visual comparisons, a Standard Entomological Metric (SEM) is used evaluate the efficacy of *wMel*-based interventions in lowering adult vector density under past and future temperature regimes: the Suppression Efficacy Score (SES). This measure is premised on the reduction of the disease-competent female population that is achieved by a given intervention.^21^ It is a single value that describes the success of vector reduction over the period of interest, here defined as one year.

We also evaluate the replacement capacity of *wMel*-based interventions under past and future climate scenarios, proposing a new SEM in service of this effort: Replacement Efficacy Score. Like the Suppression Efficacy Score, the Replacement Efficacy Score is a single value that summarizes relative success over the period of interest. However, success in this case is assessed with respect to the frequency of *Wolbachia* infection in the standing vector population over this time.^3^ A conceptual explanation of the RES score is furnished by **Figure 3**.

**Figure 3:**
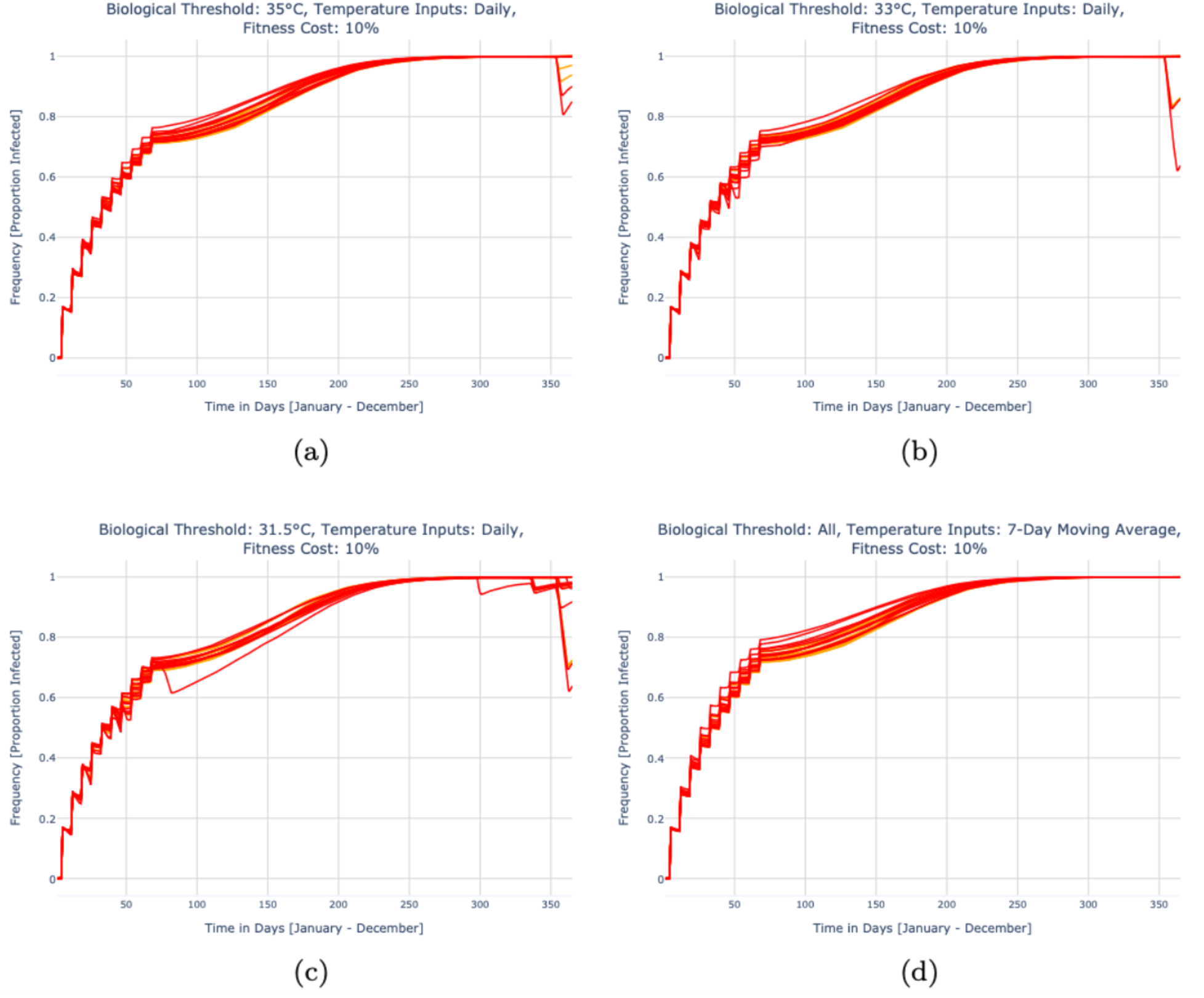
Effect of future heatwaves on the frequency of wMel-infection in an adult female Ae. aegypti population. Each line corresponds to one simulated intervention year; each color corresponds to a different temperature regime (2030 in orange, 2050 in red).

**Figure 3:**
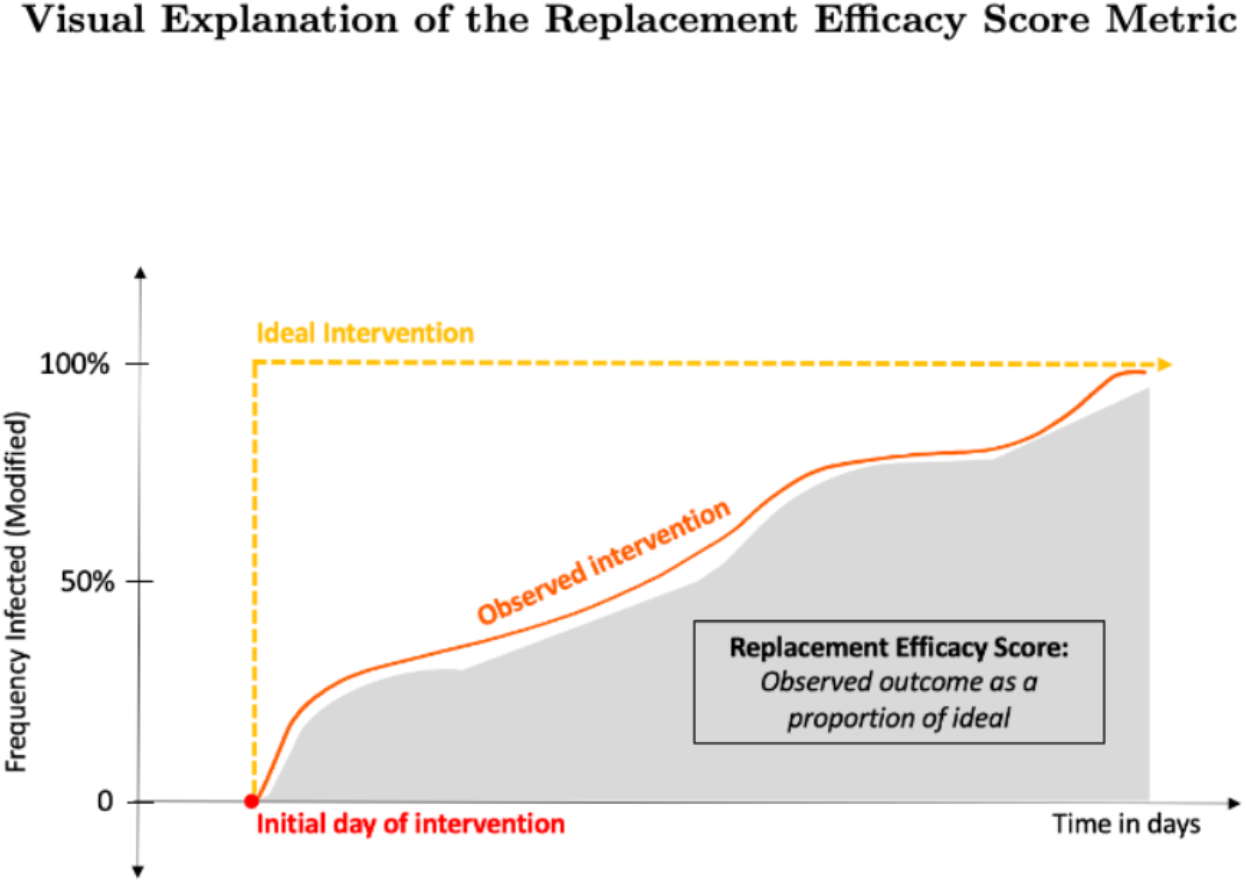
The Replacement Efficacy Score (RES) reflects the proportional comparison between the ideal and observed intervention outcomes with respect to the frequency of the modified population over the period of interest. Modification, in the case of wMel, refers to infection with the symbiont.

**Figure 4:**
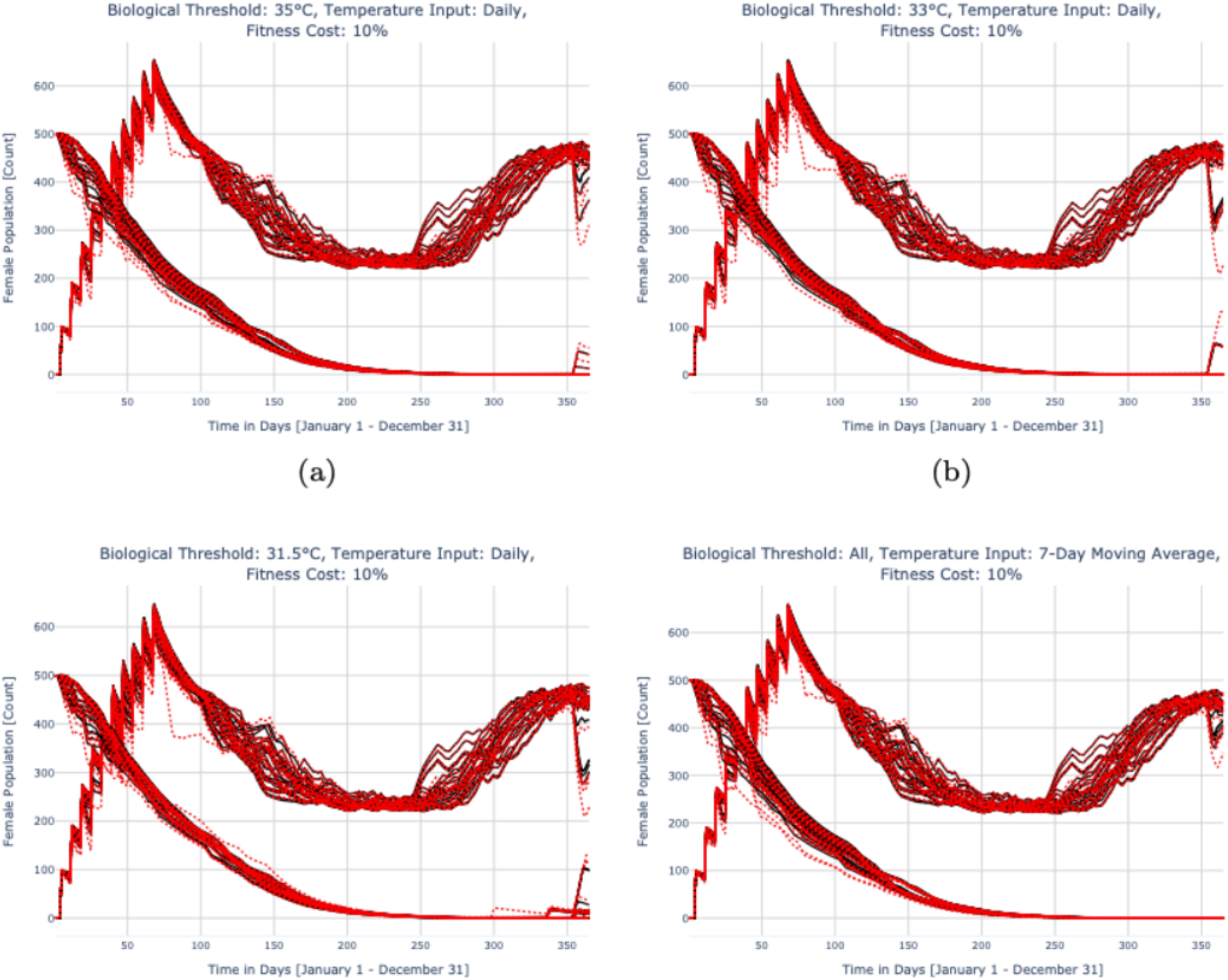
Replacement and suppression dynamics for future heatwave years (red dotted lines) and the baselines from which they are calculated (black solid lines) under both future temperature regimes (2030s 2050s). In each panel, the upper set of trajectories show the wMel-infected population and the lower set of trajectories exhibit the wildtype population.

**Figure 5:**
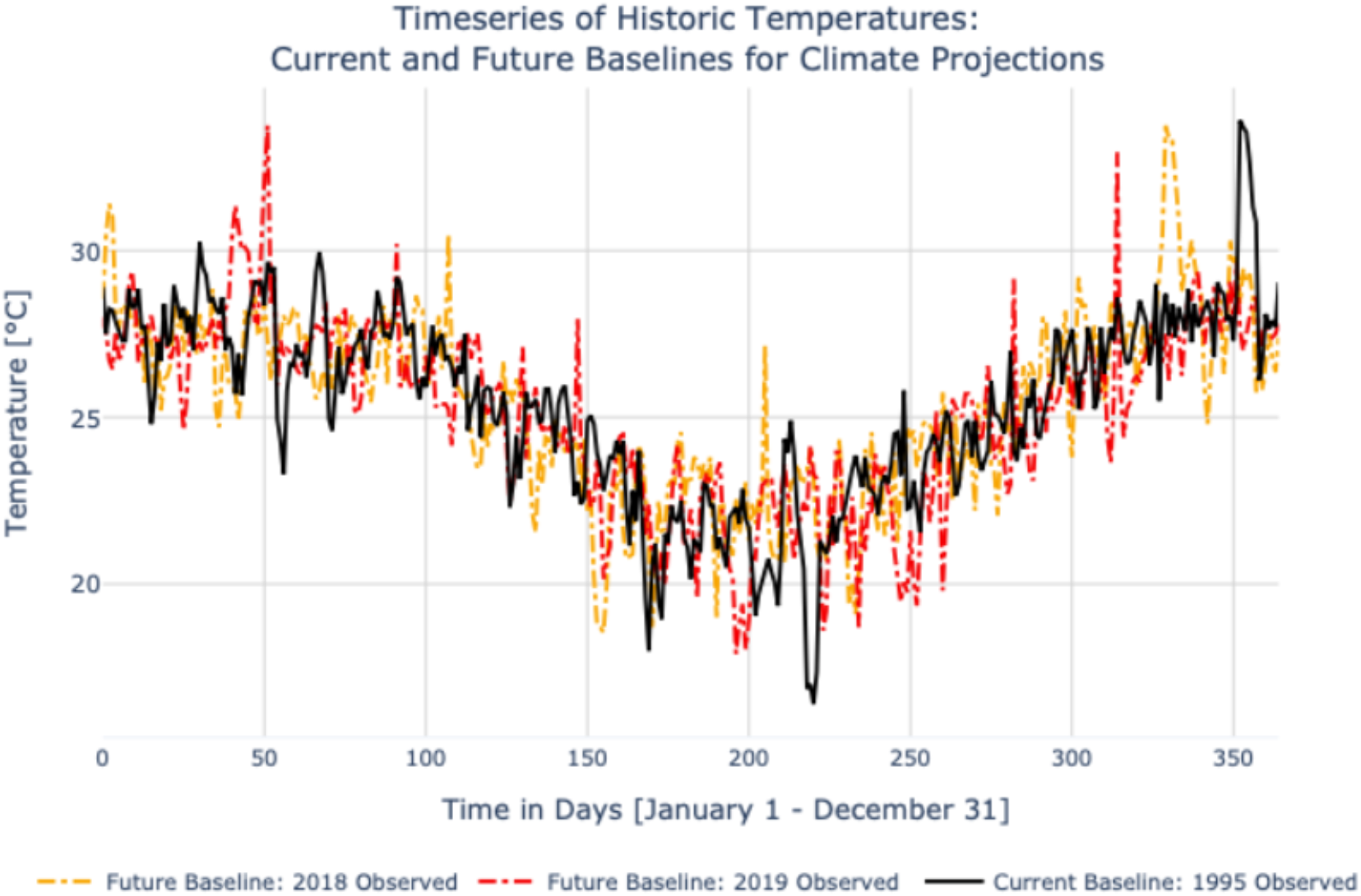
Recent years of observed temperature feature greater variability and hotter extremes than those currently used as baselines for climate projections.

**Figure 6:**
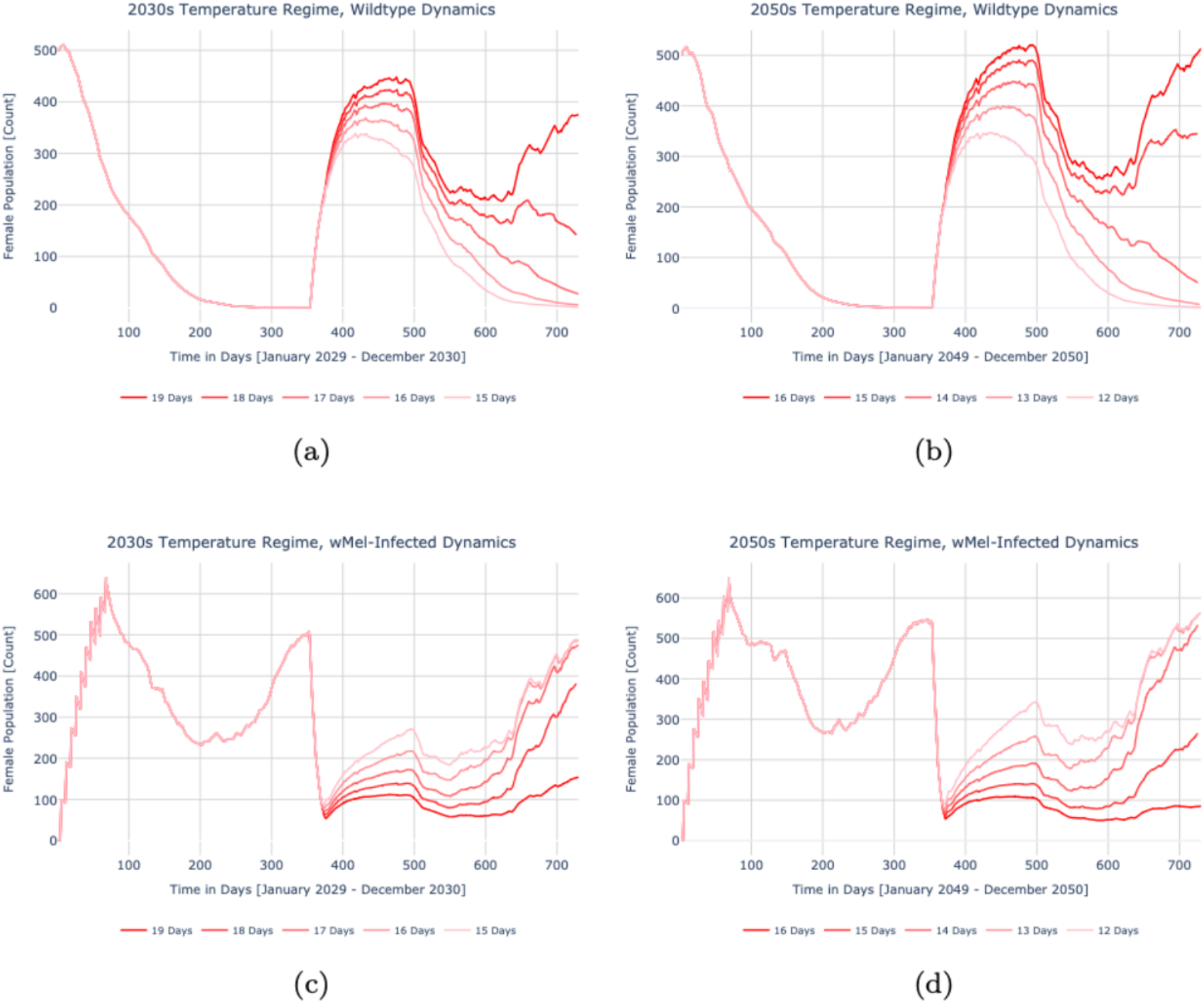
Impact of additional hot days on population dynamics in future heat­ wave years: 2030s & 2050s.

As with the SES score, in the intervention period being evaluated there are a total of *T* timesteps equal to that period’s length, τ_f_ - τ_0_, divided by the discretization, δτ. This is explained in **Equation 4**.

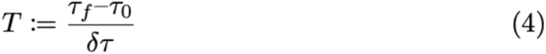

The frequency of *wMel* is estimated by calculating the proportion of the infected female population *P*_*g,t*_ and integrating the cumulative change in that population over the duration of an intervention. *A*^*F*^_*g,t*_, the area under the curve of the observed frequency trajectory, is defined in **Equation 5**.

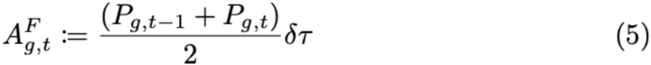

This area is then subtracted from the outcome of an idealized intervention *ρ*_g_ wherein fixation (100% infection of the standing population) is achieved immediately and for the duration of the period of interest; this yields the RES score described by **Equation 6**. Like the previously defined SEMs, indexing the RES score *R*_*g*_ and vector population change *A*^*F*^_*g,t*_ according to genotype(s) of interest *g* allows the metric to be generalized beyond *wMel*-specific applications.

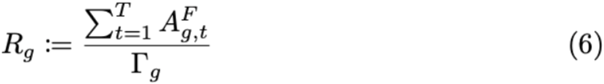

### Theoretically Testing the Thermal Limits of wMel

Using 2029 RCP 8.5 and 2049 RCP 8.5 as examples, we augmented each timeseries of temperature inputs to include, beginning on day 355, consecutive additional days of heat equivalent to the maximum value in that time series. For 2029, the maximum value was 35.21°C and for 2049 it reached 36.6°C. These temperatures are high enough to drop maternal transmission of *wMel* as well as hatch rates for *Wolbachia*-infected eggs to zero. Cytoplasmic incompatibility for crosses between wild females and *wMel*-carrying males is also eliminated. We selected the first timestep to begin appending new hot days based on the culmination of the heat surge in the 2029 example, which ended on day 354. To maintain comparability, the same timestep was chosen for the 2049 simulation. The time horizon was then extended by a second year and no additional *wMel*-infected releases were conducted.

## Footnotes

1 In 2029 RCP 4.5, the temperature reaches 35.08°C on day 353, and in RCP 8.5 it rises to 35.21°C and 35.01°C on days 353 and 354, respectively.

2 In 2049 RCP 4.5, values of 35.55°C, 35.35°C, 35.20°C are achieved on days 353, 354, and 355; in RCP 8.5 36.60°C, 36.40°C, 36.25°C, 35.35°C is reached on days 353, 354, 355, and 356.

3 The Replacement Efficacy Score is also applicable to public health intervention technologies that employ genetic modification, in which case the frequency of the refractory genotype, rather than *Wolbachia* infection, is appraised.

