## Supplementary Information for "*wMel* replacement of dengue-competent mosquitoes is robust to near-term climate change"

***Section 1: Synthesized Climate Data***

The freely available Global Historical Climatology Network (GHCN) database is a composite of climate records merged from local sources and subjected to quality assurance review; the specific records employed for this study that serve as the historical baseline were sourced from Cairns Aero (Meteorological Station ID: ASN00031011) and include date as well as daily average, maximum, and minimum temperatures for January 1, 1990 through December 31, 2019.

To produce future daily average temperatures for both RCPs, the deltas in degrees Celsius corresponding to each projected month and year from the QFCD dataset for the Queensland region were added to the historical baseline creating a single future time-series of the same length and with the same variability as the historical time-series. The future heatwave scenarios were constructed by first identifying historical heatwaves in the 1990-2019 dataset, using the Australian Bureau of Meteorology's definition: "a period of at least three days where the combined effect of excess heat and heat stress is unusual with respect to the local climate."[1,2]

We defined "unusual with respect to the local climate" as daily maximum temperatures that exceeded the historical average daily maximum temperature for the corresponding day of year by at least three degrees Celsius for at least three consecutive days. Six years within the 15-year historical baseline period containing days that met these requirements were selected: 1990, 1992, 1994, 1995, 2002, and 2005. Future heatwaves were created for the corresponding years in the future time series – specifically for 2024, 2026, 2028, 2029, 2036, and 2039 as well as 2044, 2046, 2048, 2049, 2056, and 2059. The average daily temperature for the corresponding future date of each heatwave identified within the historical record was augmented by the change in °C recorded by the "Heatwave Peak Temperature" variable in the QFCD dataset.

The duration of each heatwave identified within the historical record was also lengthened in the future time-series according to the change in days specified by the "Heatwave Duration" variable in the QFCD dataset, where each fraction of a day was rounded to the nearest full day. The additional heatwave days were given the same temperature delta in °C indicated by the "Heatwave Peak Temperature" variable, on top of the future average temperature for that date.

Finally, to reflect scientific projections of future heatwave frequency – the change in the number of heatwave days each year – every future heatwave year was augmented with an increase in the total count of heatwave days corresponding to the "Heatwave Frequency" variable in the QFCD dataset. This variable carries a distinct value for each season (wet vs. dry), each RCP scenario (4.5 vs. 8.5) and each future set of years (2030 vs. 2050). To designate a new heatwave day with consistent logic and accurate implementation while accounting for all constraints, including the requirement that "heatwaves" be defined as three consecutive days where the average daily maximum temperature meets or exceeds the average historical baseline temperature for the corresponding day of year by 3°C, a simple algorithm was developed.

For each season of each heatwave year, a running three-day average was taken of the daily maximum temperatures. These averages were sorted, greatest to least, according to the delta by which they exceeded the historical baseline for the corresponding day of year. Any day already defined as a historical heatwave day was dropped from the tally of running averages and excluded from this ordering. The N hottest averages, where N = (Heatwave Frequency)/3, that were at minimum four days apart from each other, were designated as new future heatwave days. The average daily temperature of the newly selected days, together with the two adjacent days that together informed the qualifying three-day running averages, was then augmented by the value of the "Heatwave Peak Temperature" variable.

**Figure 1** features a temporal subset of four timeseries created using this methodology for the years 2029 and 2049 from the historic baseline of 1995. It illustrates the differential between 2030 and 2050 temperature regimes for both RCP Scenarios 4.5 and 8.5 in the case of heatwaves as well as average temperature.

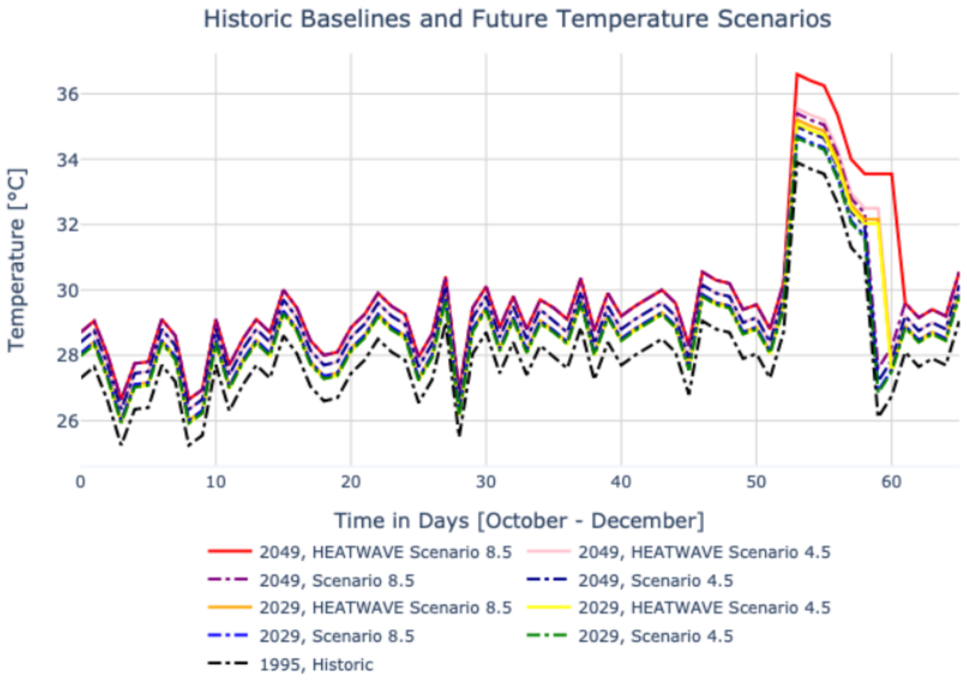


Figure 1: Temporal subset of example temperature time series developed using RCP 4.5 and 8.5 scenarios.

***Section 2: Mosquito Population Model***

The model equations representing the change in numbers of *Ae. aegypti* for the various life stages within a single geographic node are as follows. They assume a timestep of one day; all simulations run using them in this work are on a time horizon of one year unless otherwise noted.

$$\omega_g = \sum_{i=1}^{N} \beta_g \sigma_g \left( \mathbf{\Gamma}_g \odot \mathbf{T}_g \right)_i F_i \qquad \forall g \qquad (1a)$$

$$\frac{dE_{g,1}}{dt} = \omega_g - E_{g,1}(\mu_E + q_E n_E) \qquad \forall g \qquad (1b)$$

$$\frac{dE_{g,i}}{dt} = E_{g,i-1} q_E n_E - E_{g,i}(\mu_E + q_E n_E) \qquad \forall g, i = 2 \ldots n_E \qquad (1c)$$

$$\frac{dL_{g,1}}{dt} = E_{g,n_E} q_E n_E - L_{g,1}(\mu_L d + q_L n_L) \qquad \forall g \qquad (1d)$$

$$\frac{dL_{g,i}}{dt} = L_{g,i-1} q_L n_L - L_{g,i}(\mu_L d + q_L n_L) \qquad \forall g, i = 2 \ldots n_L \qquad (1e)$$

$$\frac{dP_{g,1}}{dt} = L_{g,n_L} q_L n_L - P_{g,1}(\mu_P + q_P n_P) \qquad \forall g \qquad (1f)$$

$$\frac{dP_{g,i}}{dt} = P_{g,i-1} q_P n_P - P_{g,i}(\mu_P + q_P n_P) \qquad \forall g, i = 2 \ldots n_P \qquad (1g)$$

$$\frac{dm_g}{dt} = P_{g,n_P} q_P n_P (1 - \theta_g) - m_g \mu_m \qquad \forall g \qquad (1h)$$

$$X_g = P_{g,n_P} q_P n_P \theta_g \frac{m_g \eta_g}{\sum_{k=1}^{N} m_k \eta_k} \qquad \forall g \qquad (1i)$$

$$\frac{dF_{g,i}}{dt} = X_{g,i} - F_{g,i} \mu_F \qquad \forall g, i \qquad (1j)$$

Here, the number of individuals in juvenile stages of egg, larva, and pupae are represented by variables $E$, $L$, and $P$ and distinguished by genotype $g$. Adult stages male, $m$,[1] and female, $F$, are assumed to mate once immediately upon emergence from the pupal stage. Mated females are represented by $X$.

Wildtype mortality rates $\mu$ and development rates $q$ are dynamically calculated according to temperature using the functional forms specified by Rossi et al (2014).[3] Fitness cost is implemented as a 0%, 10%, or 20% increase in $\mu$ of both adult stages. *Wolbachia*-infected mortality rates $\mu$, inheritance probabilities $\Gamma_g$, and survival probabilities $T_g$ are dynamically calculated according to temperature as described in the Methods section of this work.

Logistic density dependence $d$ is implemented in the larval stage $L$. The number of eggs laid per genotype is represented by $\omega_g$, while $\beta_g$ and $\sigma_g$ are female and male fecundity parameters and $\Gamma_g$ and $T_g$ convey the inheritance and survival probability of the specified genotypes, respectively. The parameterization of $\Gamma_g$ follows Sánchez et al (2020)[4] while the formulation of Equations 1a and 1i follow Sánchez et al (2020).[5]

[1] Referred to using a lowercase letter because, unlike the juvenile or female stages, this stage is implemented as a vector rather than a matrix.

The GeneDrive.jl software extended to implement this work is located on GitHub at https://github.com/vnvasquez/GeneDrive.jl; all code adaptations required to reproduce the specific experiments in this paper are included in the https://github.com/vnvasquez/WolbachiaClimatePaper repository. The raw data outputs featuring the population dynamics that resulted from all model runs is stored on Figshare: 10.6084/m9.figshare.20722270, 10.6084/m9.figshare.20721928, 10.6084/m9.figshare.20719234, 10.6084/m9.figshare.20716117, 10.6084/m9.figshare.20607765.

***Section 3: Additional Results and Sensitivity Analyses***

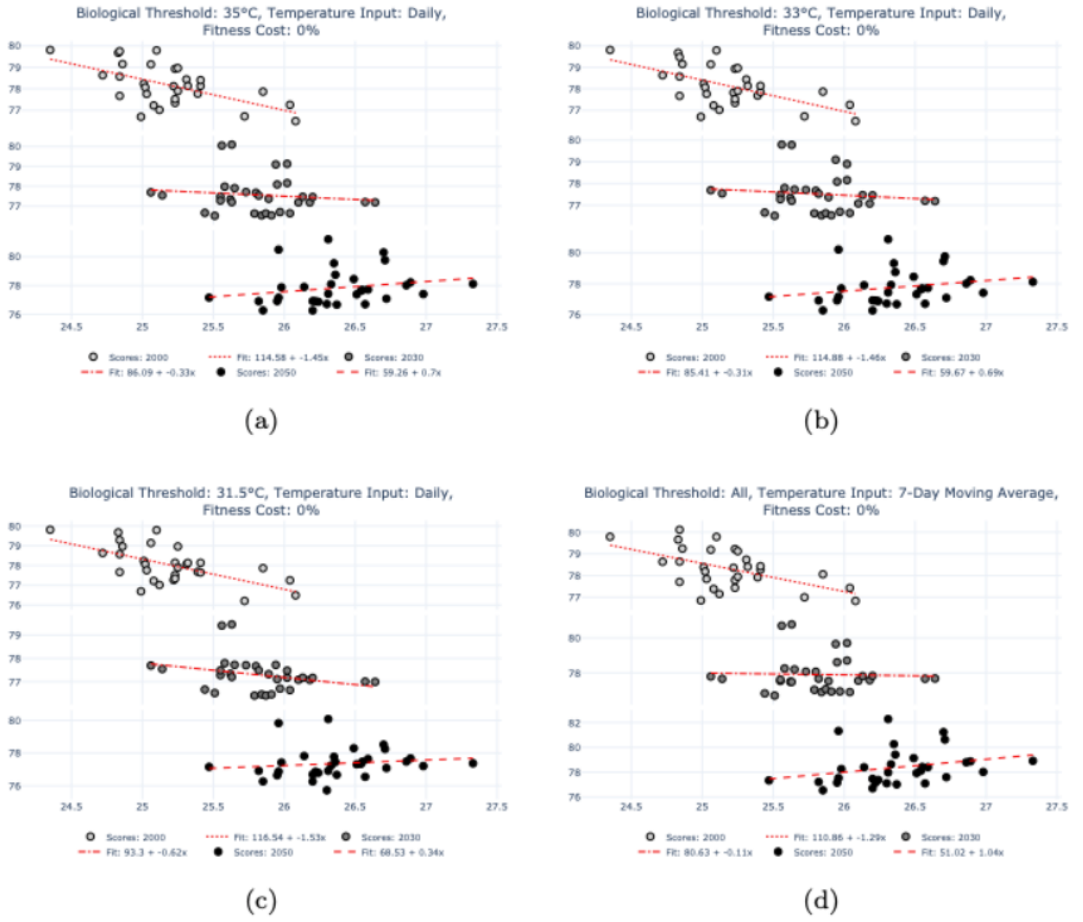


Figure 2: Increases in average temperature impact the suppression efficacy of Wolbachia-based interventions. Here, Suppression Efficacy Scores (SES) at 0% fitness cost.

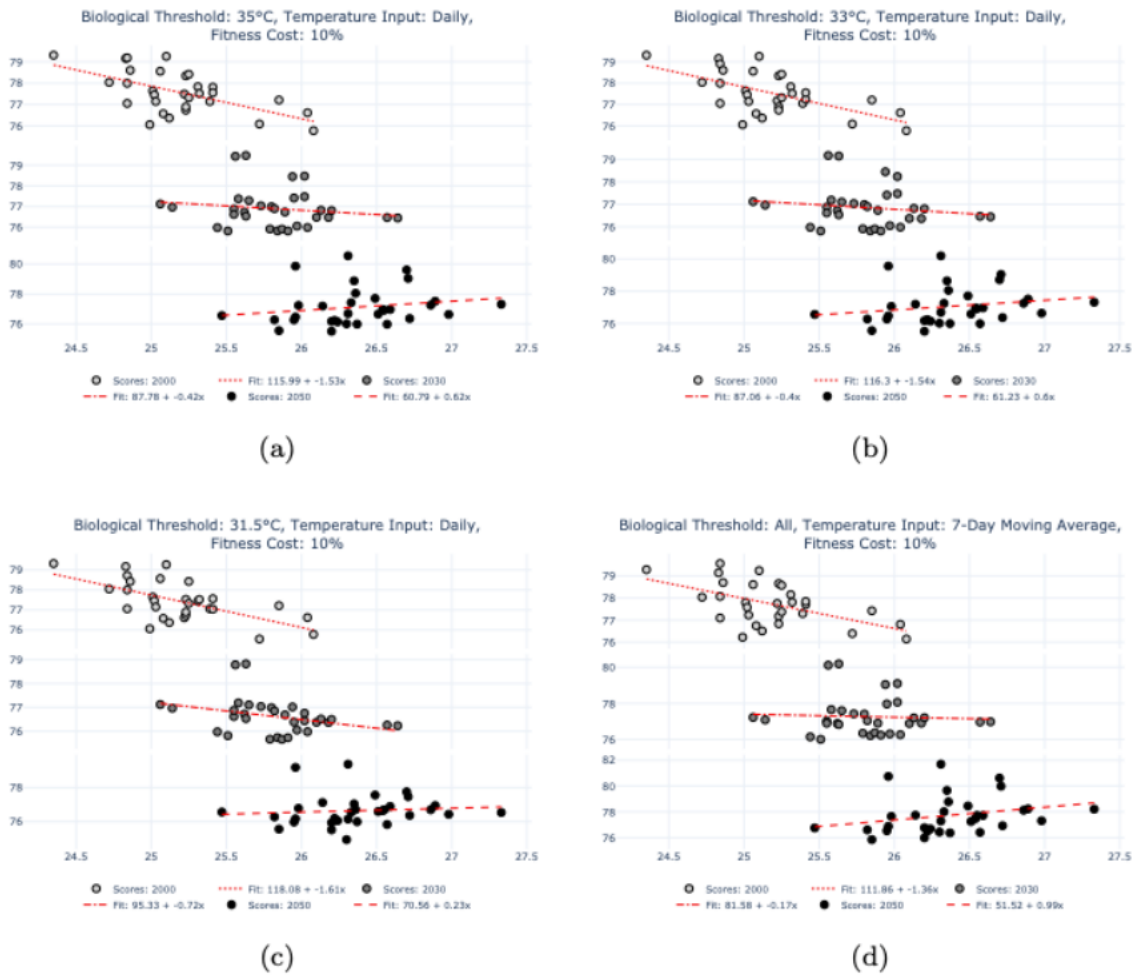


Figure 3: Effect of rising average temperature on Suppression Efficacy Scores (SES) at 10% fitness cost.

**Supplementary Information**: *wMel* replacement of dengue-competent mosquitoes is robust to near-term climate change (Vásquez et al 2022)

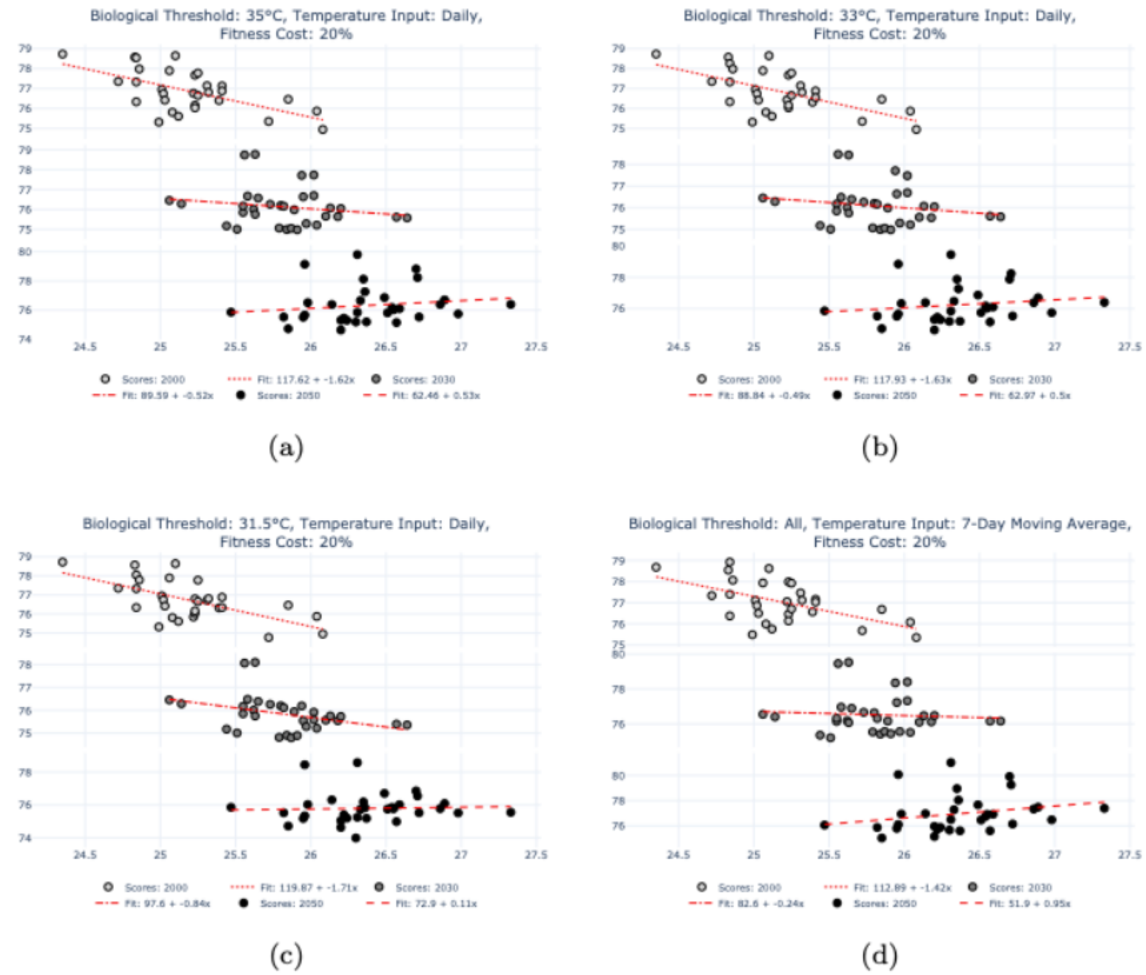


Figure 4: Effect of rising average temperature on Suppression Efficacy Scores (SES) at 20% fitness cost.

**Supplementary Information**: *wMel* replacement of dengue-competent mosquitoes is robust to near-term climate change (Vásquez et al 2022)

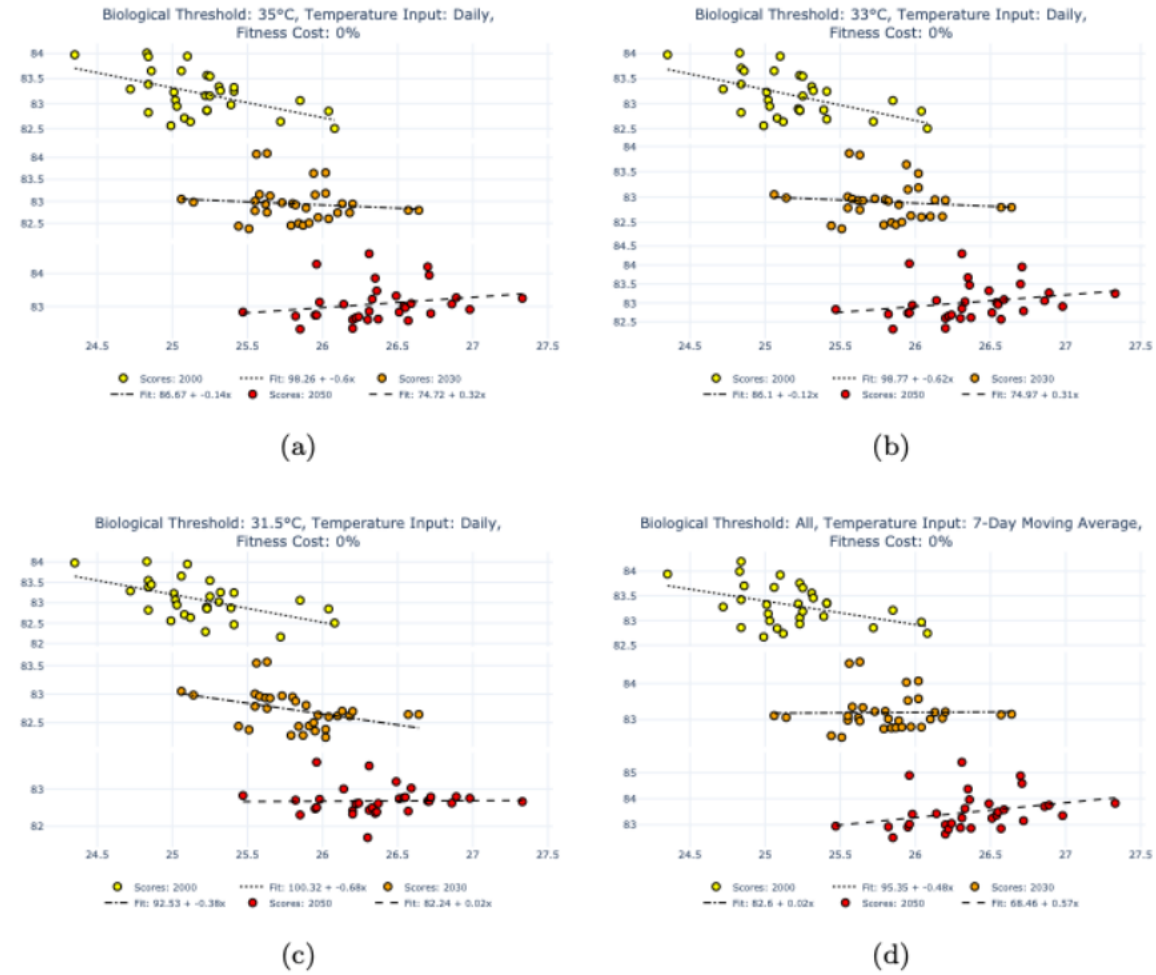


Figure 5: Effect of rising average temperature on Replacement Efficacy Scores (RES) at 0% fitness cost.

**Supplementary Information**: *wMel* replacement of dengue-competent mosquitoes is robust to near-term climate change (Vásquez et al 2022)

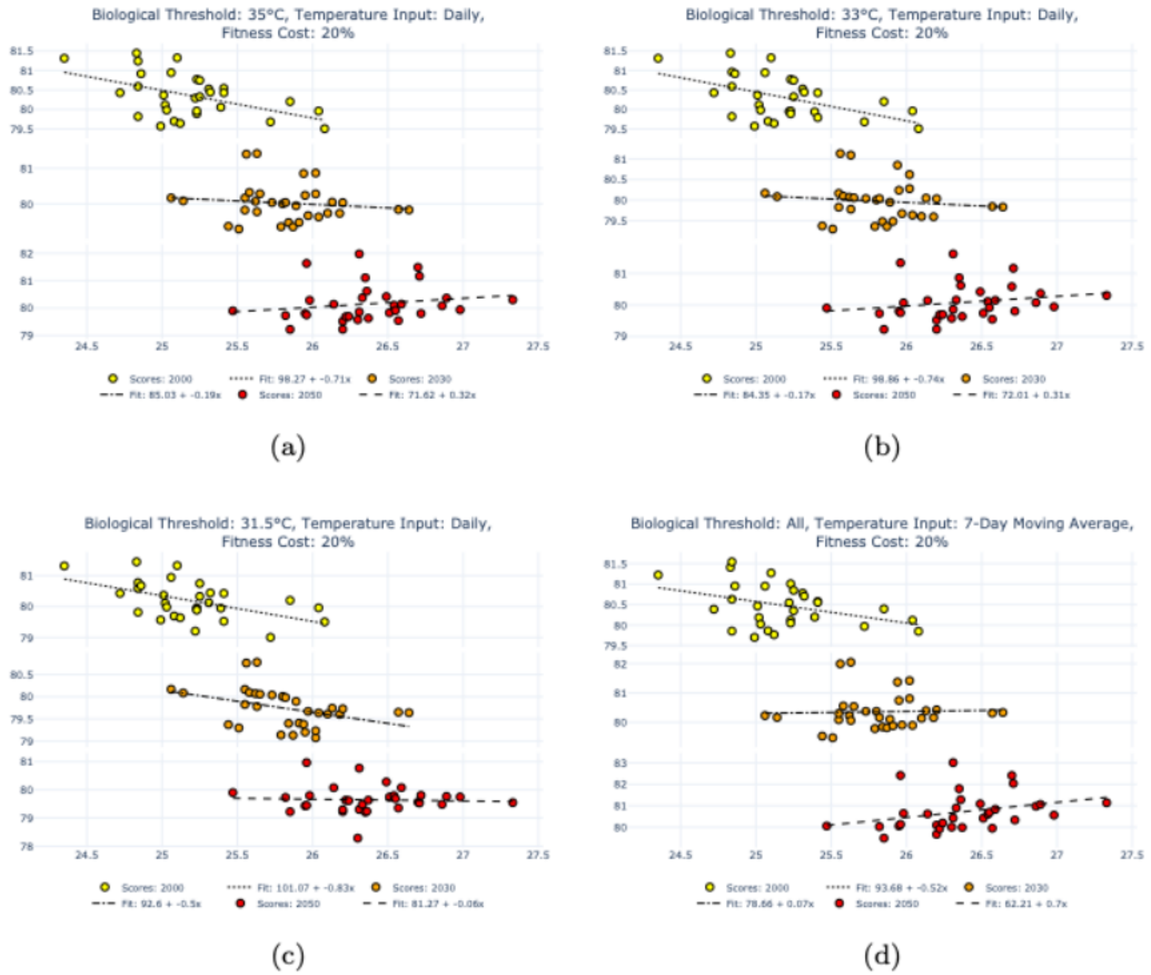


Figure 6: Effect of rising average temperature on Replacement Efficacy Scores (RES) at 20% fitness cost.

**Supplementary Information**: *wMel* replacement of dengue-competent mosquitoes is robust to near-term climate change (Vásquez et al 2022)

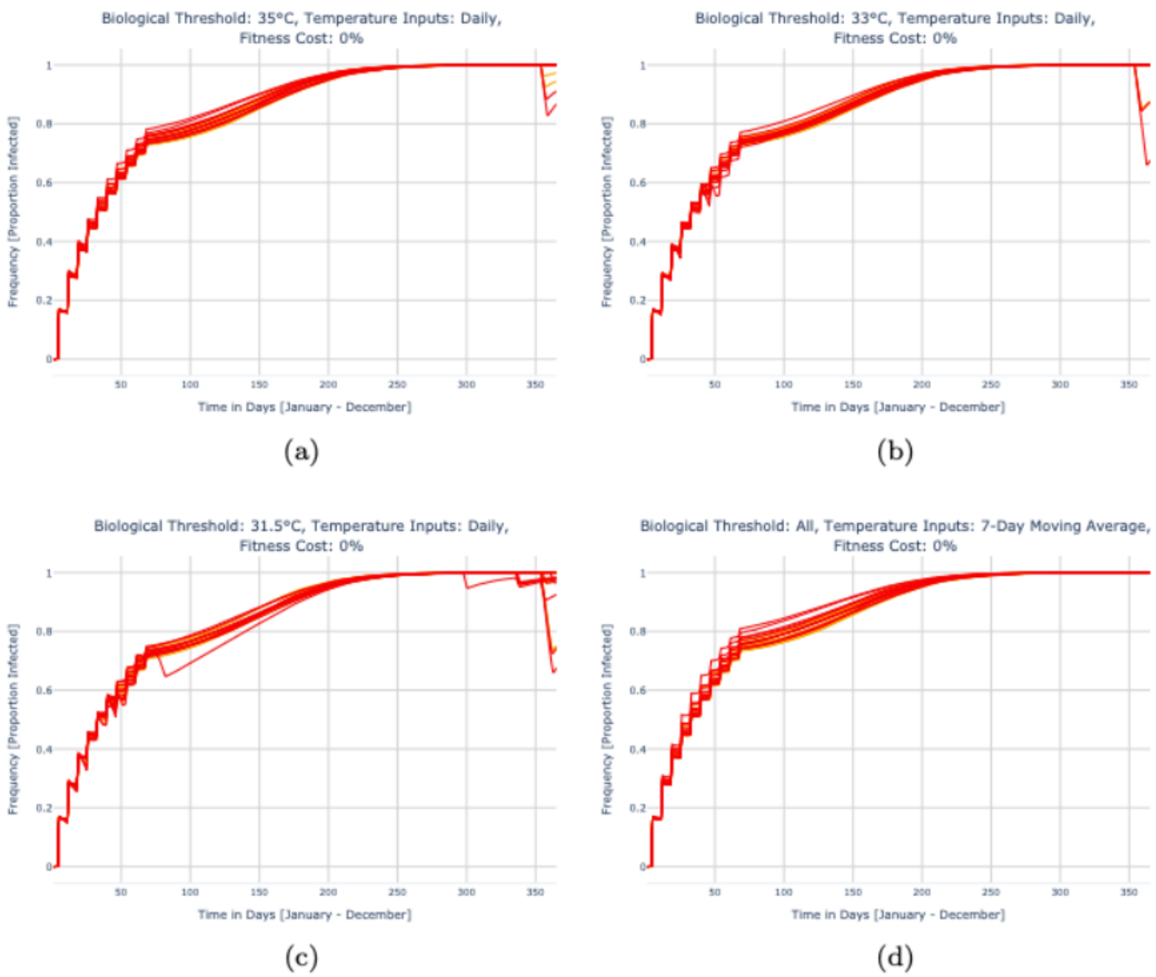


Figure 7: Effect of future heatwaves on the frequency of wMel infection in an adult female Ae. aegypti population assuming 0% fitness cost.

**Supplementary Information**: *wMel* replacement of dengue-competent mosquitoes is robust to near-term climate change (Vásquez et al 2022)

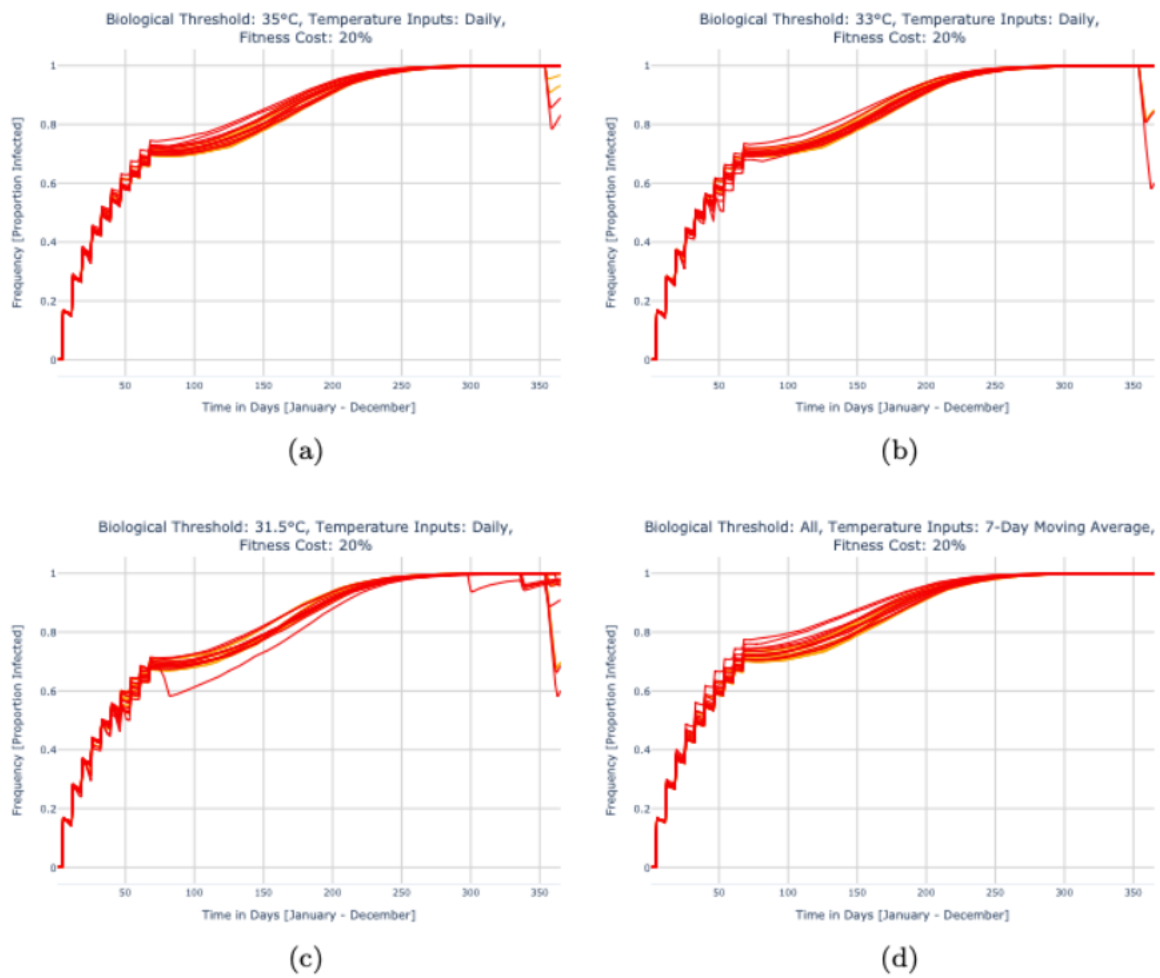


Figure 8: Effect of future heatwaves on wMel infection frequency assuming 20% fitness cost.
